# Human IL-2 receptor β mutations associated with defects in immunity and peripheral tolerance

**DOI:** 10.1101/476697

**Authors:** Zinan Zhang, Florian Gothe, Perrine Pennamen, John James, David MacDonald, Carlos P. Mata, Yorgo Modis, Meghan Acres, Wolfram Haller, Claire Bowen, Rainer Doffinger, Jan Sinclair, Shannon Brothers, Anas Alazami, Yu Zhang, Helen Matthews, Sophie Naudion, Fanny Pelluard, Yasuhiro Yamazaki, Luigi Notarangelo, Hamoud Almousa, James Thaventhiran, Karin R. Engelhardt, Sophie Hambleton, Caroline Rooryck, Ken Smith, Michael J. Lenardo

**Author notes:** co-first authors (Zinan Zhang, Florian Gothe, Perrine Pennamen). co-senior authors (Sophie Hambleton, Caroline Rooryck, Ken Smith, Michael J. Lenardo).

## Abstract

Interleukin-2, which conveys essential signals for effective immunity and immunological tolerance, operates through a heterotrimeric receptor comprised of α, β and γ chains. Genetic deficiency of the α or γ chain causes debilitating disease. Here we identify human interleukin-2 receptor (IL-2R) β chain (CD122) gene defects as a cause of life-threatening dysregulation of immunity and peripheral tolerance. We report homozygous mutations in the human *IL-2Rβ* gene from four consanguineous families, comprising either recessive missense mutations in five children or an early stop-gain mutation in two deceased fetuses and a premature neonate. All patients surviving to childhood presented with autoantibodies, hypergammaglobulinemia, bowel inflammation, and dermatological abnormalities, as well as cytomegalovirus disease in most cases. Patient T lymphocytes lacked surface expression of IL-2Rβ and were unable to normally respond to high-dose IL-2 stimulation. By contrast, patient natural killer (NK) cells retained partial IL-2Rβ expression and cytotoxic function. IL-2Rβ loss of function was recapitulated in a recombinant system, in which endoplasmic reticulum sequestration was revealed as the mechanism by which certain missense mutations cause disease. Hematopoietic stem cell transplant resulted in resolution of clinical symptoms in one patient. The hypomorphic nature of this disease highlights the significance of variable IL-2Rβ expression in different lymphocyte subsets as a means of modulating immune function. Insights from these patients can inform the development of IL-2-based therapeutics for immunological diseases and cancer.

## Introduction

The interleukin-2 receptor (IL-2R) complex plays a central role in control of the immune response by integrating signals from the key cytokines IL-2 and IL-15. Three distinct receptors for IL-2 are generated by combinations of three IL-2R subunits: IL-2Rα(CD25), IL-2Rβ (CD122) and IL-2Rγ(CD132) – the latter, known as the common γchain, is also necessary for signaling by IL-4, 7, 9, 15 and 21. All three chains combine to form the high affinity IL-2R, which is constitutively expressed on CD4+ regulatory T cells (T_regs_), and induced upon activation of CD4 and CD8 T cells, B cells and some myeloid-derived subsets (Liao et al. 2013; Busse et al. 2010). A second receptor, which binds IL-15 and IL-2 with intermediate affinity, is comprised of only the IL-2Rβ and IL-2Rγ subunits; it is constitutively expressed on resting CD8+ T cells and natural killer (NK) cells. The α subunit alone is a low affinity receptor. Upon ligand binding, the IL-2Rβ and IL-2Rγ subunits undergo tyrosine phosphorylation which, in turn, induces the phosphorylation of the associated Janus tyrosine kinases (JAK) 1 and 3, that phosphorylate the signal transducer and activator of transcription 5 (STAT5) transcription factor (Waldmann et al. 2006). STAT5, once dimerized and translocated to the nucleus, induces a pro-survival and proliferation transcription program. Interleukin-2 (IL-2) is primarily produced by CD4+ T helper cells following T cell receptor (TCR) engagement with costimulation (Boyman et al. 2012). It potently stimulates T cell proliferation, differentiation (promoting Th1, Th2, and Th9, and suppressing Th17 polarization) and cytolytic effector activity. It also plays a key role in peripheral tolerance by promoting the generation and maintenance of regulatory T cells (T_reg_) and antigen-specific peripheral T cell clonal deletion (Hatakeyama et al. 1989; Takeshita et al 1992; Lenardo, 1991). CD25 deficient mice demonstrate grossly normal early B and T cell development, but lymphadenopathy and impaired T cell activation and clonal deletion. As they age, these mice develop autoimmune and inflammatory disease (e.g. hemolytic anemia and inflammatory enteropathy) (Willerford et al. 1995). Humans with CD25 deficiency have a similar phenotype, developing prominent autoimmune disease with less consistent evidence of immunodeficiency, and resembling patients with IPEX, due to FOXP3 deficiency, thus indicating the impact of loss of the high-affinity IL-2 receptor can be ascribed to a loss of peripheral tolerance (Scharfe et al. 1997, Caudy et al. 2007).

The role in immunity of IL-2Rβ is less well understood, and no monogenic cause of human IL-2Rβ deficiency has yet been described. IL-2Rβ-deficient mice had severe autoimmunity and diminished cytolytic effector function, with splenomegaly, lymphadenopathy, elevated IgG1 and IgE levels, and ANA and anti-dsDNA autoantibodies (Suzuki et al. 1995). They succumbed to autoimmunity around 12 weeks and this can be reversed by the adoptive transfer of T_regs_ (Malek et al. 2002). Despite showing evidence of activation (e.g. increased CD69 and CD25), the T cells of IL-2Rβ-deficient mice failed to respond to stimuli including IL-2, PMA, and ionomycin, and had diminished CD8 T+ cell cytolytic activity when re-challenged (Suzuki et al. 1995). This, plus the observation that they have reduced NK cell numbers (Suzuki et al. 1997), implies that IL-2Rβ deficiency in mice could produce susceptibility to infection in addition to T cell activation and autoimmunity. IL-2Rβ-mediated signaling is implicated in pathways known to be important in human autoimmune disease, and loci containing it have been associated with asthma and juvenile-onset arthritis in genome-wide association studies (Moffatt et al. 2010, Hinks et al. 2013). Moreover, high-dose IL-2 therapy is approved for use in renal cell carcinoma and malignant melanoma and encouraging early phase studies of low-dose IL-2 in Type-1 diabetes, graft-versus-host disease and systemic lupus erythematosus have led to over 14 on-going phase 2 and 3 trials (Ahmadzadeh et al. 2006; Ye et al. 2018). It will thus be important to understand the biology of IL-2Rβ, and the impact of IL-2Rβ deficiency, in humans. To this end, we describe human homozygous recessive IL-2Rβ deficiency in four consanguineous families with 8 affected individuals, which was associated with autoimmunity and immunodeficiency.

## Results

### Clinical phenotype and genotype of patients in a combined immunodeficiency/autoimmunity cohort

We investigated the medical records and clinical data of eight affected individuals from four consanguineous families with South Asian, Middle Eastern, and Eastern European origins, who now reside in countries on three different continents. All the patients have a history of severe immunodeficiency and autoimmunity (Figure 1 and Supplementary table 1). Kindred A includes a six-year-old boy (A1) and his two-year-old sister (A2) born to first cousin Pakistani parents (Figure 1A). A1 was initially hospitalized at the age of two for thyrotoxicosis secondary to Graves’ disease and A2 was hospitalized at the age of six months for failure to thrive and persistent diarrhea (Supplementary table 1). Since their initial hospital course, A1 has developed severe gastroenteritis and dermatitis and A2 has had pulmonary, gastrointestinal, and urinary infections as well as ANCA+ vasculitis. Patient A1 has improved with rituximab treatment but continues to be intermittently ill. Patient A2 received an allogeneic hematopoietic stem cell transplant (HSCT) and has recovered with no sequelae. Kindred B consists of a girl (B1) born to related parents of South Asian origin. B1 initially presented in a collapsed state with severe diarrhea at the age of 4 weeks and was found to have enteropathy, dermatitis, and later CMV viremia with hepatitis. She improved with immunosuppression but ultimately succumbed to probable CMV pneumonitis and respiratory failure after HSCT. Kindred C includes a boy (C1) and his first female cousin (C2) born to consanguineous Saudi Arabian parents. C1 presented with suppurative ear infections at the age of 6 months. C2 presented with chest and ear infections and diarrhea at the age of 2 months. Subsequently, both suffered recurrent otitis, severe dermatitis, CMV viremia and food allergies. C1 and C2 died from probable CMV pneumonitis at the age of 3 years old and 18 months old, respectively. Kindred D consists of a premature female neonate (D1) and two fetuses (D2, D3) conceived by a Romany family living in France. D1, D2, and D3 were found to have intra-uterine growth retardation and fetal immobilism; skin-like floating membranes were also present in the amniotic fluid in all three cases. D1 was delivered pre-maturely by emergency Cesarean section at 31 weeks old, but the female neonate died two hours later due to diaphragmatic immobility. D2 and D3 pregnancies were terminated due to fetal abnormalities at 25 weeks and 30 weeks, respectively. In summary, all the patients had recurrent infections as well as autoimmune disease leading to early death in most cases.

**Table 1.**
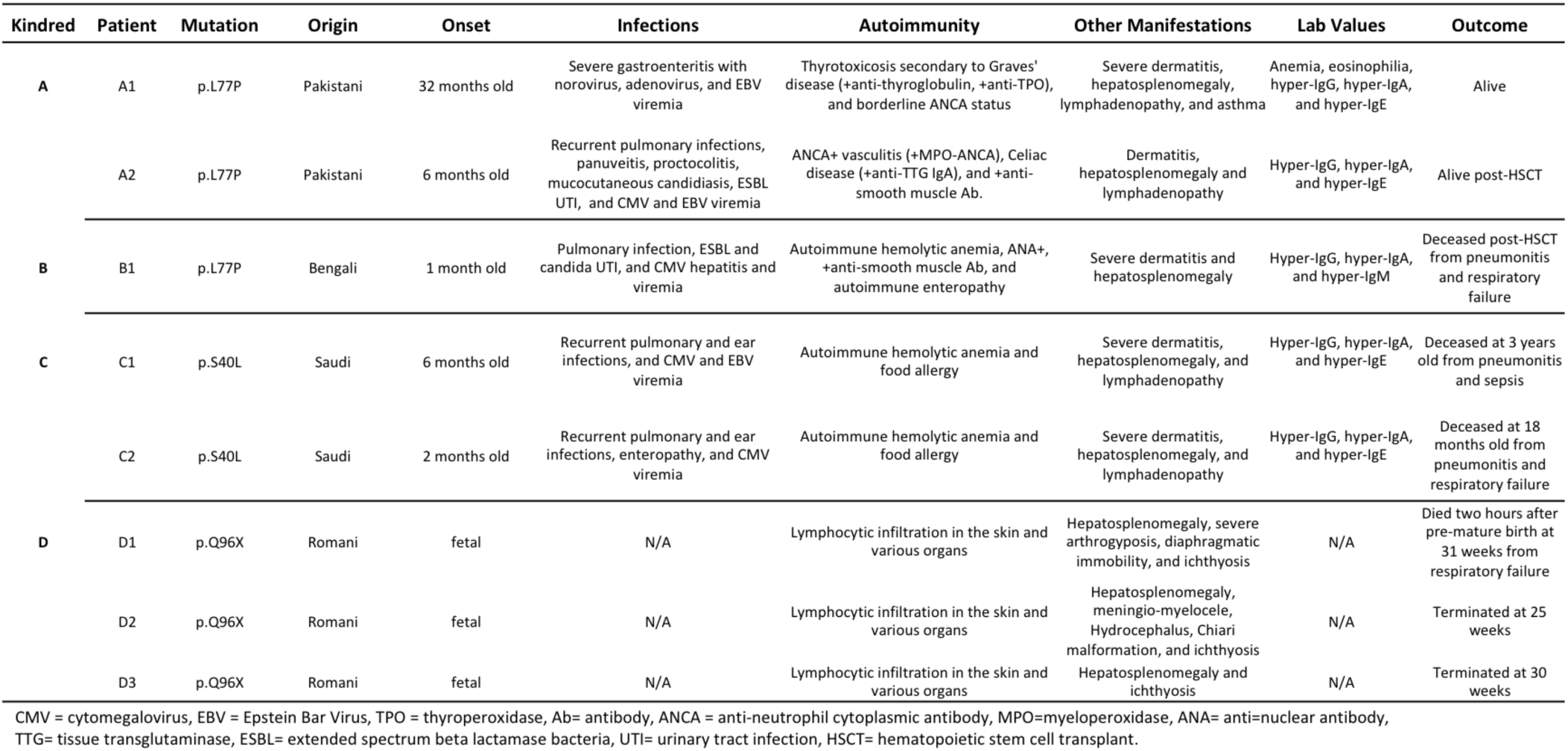
Patient mutations and clinical manifestations.

**Figure 1.**
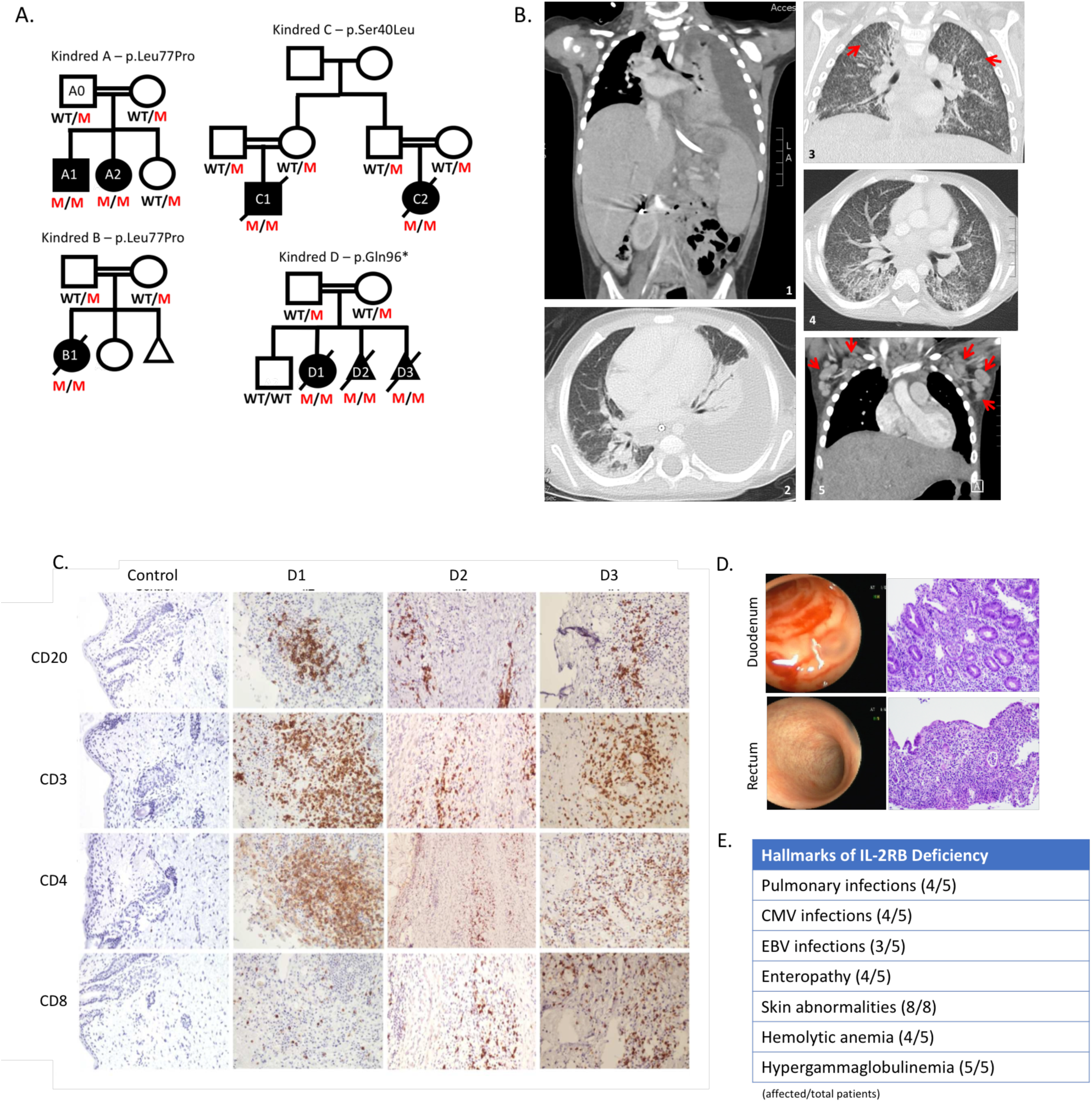
Genetic and clinical features of the disease cohort. A. Four consanguineous pedigrees of eight affected individuals (A1-D3) with three different homozygous recessive mutations. B. Radiographic evidence for pulmonary disease in Kindred A. Panels 1 and 2 show a left pleural effusion. Hepatosplenomegaly can also be seen in Panel 1. Panels 3 and 4 show numerous small pulmonary nodules and tree-in-bud changes suggestive of pneumonia. Red arrows highlight 2 small lung nodules. Panel 5 shows enlarged axillary lymph nodes (red arrows). C. Immunohistochemistry of fetal skin from kindred D, patients D1, D2, and D3 stained for the lymphocyte markers as indicated. D. Immunohistochemistry of duodenum (top) and rectum (bottom) biopsies and corresponding endoscopy images from Kindred B. E. Summary of clinical hallmarks of IL-2Rβ deficiency in the five pediatric patients. Skin abnormalities were observed in the individuals in kindred D in addition to the pediatric patients (8 total).

Immune dysregulation was a key shared feature across these four kindreds, manifested as enteropathy, dermatitis, autoimmune hemolytic anemia, and hypergammaglobulinemia (Figure 1). All patients who survived beyond the neonatal period also had recurrent infections, including defective immune suppression of herpesviruses (CMV or EBV viremia in all; CMV disease in 4 of 5; Supplementary Table 1). Chest radiographs of patient A2 revealed a pleural effusion and numerous small pulmonary nodules and tree-in-bud opacities suggestive of CMV pneumonia in the context of CMV viremia (Figure 1B). CT imaging also revealed hepatosplenomegaly and marked lymphadenopathy in A2 (Figure 1B); a clinical feature that was noted in all five patients.

Skin abnormalities are a key hallmark of this disease. A1, A2, B1, C1, and C2 all had severe dermatitis and D1, D2, and D3 had ichthyosis and significant infiltration of B and T lymphocytes on skin immunohistochemistry (Figure 1C). Four out of the five children have also had severe diarrhea and infectious/autoimmune enteropathy. Endoscopy of patient B1 showed villous atrophy and gastrointestinal biopsies revealed chronic inflammatory infiltration of the duodenum and rectum (Figure 1D). Additional hallmarks of disease include: autoimmune hemolytic anemia (4/5 patients) and hypergammaglobulinemia (5/5 patients) comprising predominantly class-switched isotypes: IgA, IgG, and IgE (Figure 1E and Supplementary Table 2). Overall, CD4 cell numbers were normal but two of the patients had low CD8 T cell counts, while NK numbers were increased (Supplementary Table 2).

**Table 2.** Lab values and absolute cell counts.

| Lab Values | A1 | A2 | B1 | C1 | C2 | Reference |
| --- | --- | --- | --- | --- | --- | --- |
| CMV | Negative | 37407 | Positive | 10000 | 2340 | Negative (0 copies/ml) |
| EBV | 2664 | 80254 | Negative | 157180 | Negative | Negative (0 copies/ml) |
| IgM | 0.34 | 1.0 | 6.6 | 0.7 | 0.86 | (0.37-1.84 g/L) |
| IgA | 4 | 8.4 | 1.55 | 4.96 | 3.86 | (0.2-1.2 g/L) |
| IgG | 22.3 | 23 | 26.9 | 22.2 | 12.3 | (2.5-9.1 g/L) |
| IgE | 12180 | 3031 | 20 | 24990 | 8370 | (1.6-30kU/L) |
| Anemia | Coombs+ | Negative | Coombs + | Coombs + | Coombs + | Negative |
| Absolute Cell Counts |  |  |  |  |  |  |
| CD3+ | 1272 | 3897 | 3454 | 1583 | 4098 | 1700-1900 cells/ul |
| CD3+/CD4+ | 954 | 2328 | 1436 | 950 | 2466 | 800-1700 cells/ul |
| CD3+/CD8+ | 291 | 1429 | 1906 | 361 | 1128 | 700-1000 cells/ul |
| CD19+ | 859 | 1274 | 533 | 728 | 1760 | 400-800 cells/ul |
| CD16+/CD56+ | 321 | 1728 | 574 | 538 | 503 | 200-400 cells/ul |

### Identification of protein-coding mutations in the gene encoding IL-2Rβ (CD122)

Because of the early onset of disease in consanguineous families, we sought a genetic cause for the disease using whole exome DNA sequence analysis of the 4 kindreds. We identified three different *IL-2Rβ* gene mutations (Figure 2A). For Kindreds A and B, the *IL-2Rβ* chr22: g.37538526 A>G (p.Leu77Pro) missense variant was prioritized. The mutation occurs in exon 4 (out of 10) and is not found in dbSNP, ESP, or ExAC databases, but has a MAF of 0.00001218 in gnomAD. The p.Leu77Pro mutation introduces a restrictive proline-proline motif in the extracellular D1 domain of IL-2Rβ (Figure 2B). For Kindred C, the g.37539634 C>T (p.Ser40Leu) missense variant was prioritized and not found in any databases. The mutation appears to be located at the interface of IL-2Rβ and IL-2 (Figure 2B). For Kindred D, a g.37537259 G>A (p.Gln96*) stop-gain mutation was identified and also not found in any databases. This mutation would lead to significant truncation of the 552 amino acid protein. Due to the predicted deleterious nature of these variants, their segregation with disease and the similarity in phenotype with a mouse knock-out model, IL2RB represented an attractive candidate disease gene. Other prioritization criteria that were taken into consideration include: CADD score, quality of reads, GERP conservation score, co-segregation of alleles, SIFT score, PolyPhen2 score, tissue specific expression levels, structural modeling, and primary literature reviews leading to the conclusion that these variants were likely responsible for the disease observed in the respective patients in this cohort.

**Figure 2.**
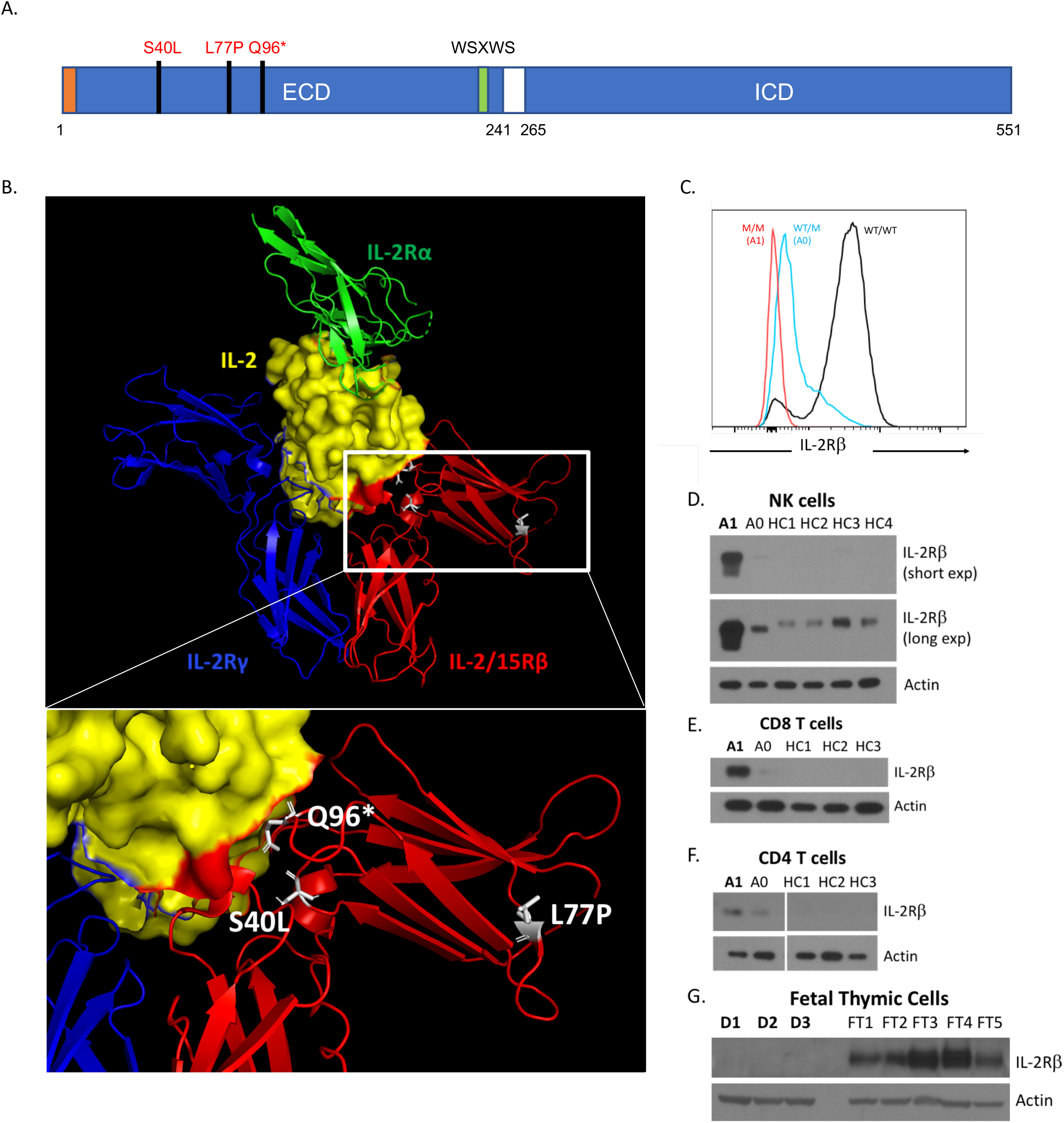
IL-2Rβ coding mutations causes IL-2Rβ surface receptor deficiency. A. Schematic of intracellular (ICD) and extracellular domains (ECD) of the IL-2Rβ protein depicting the location of the three mutations in the ECD. The signal peptide is highlighted in orange and the canonical WSXWS motif is highlighted in green. B. Crystal structure of IL-2:IL-2R complex with the expanded view showing the position of the three mutations in white: L77P, S40L, and Q96*; (modified from PDB 2B5I, Wang et al. 2005). Red: IL-2/15Rβ, blue: IL-2Rγ, green: IL-2Rα, and yellow: IL-2 with IL-2Rβ interface colored in red. C. Histogram of IL-2Rβ surface expression in NK cells (CD3^−^ CD56^+^) (red = homozygous affected A1, blue = heterozygote healthy A0, black=healthy control). D. Western blot of FACS-sorted CD3^−^ CD56^+^ NK cells from A1, heterozygote parent (A0), and four healthy controls (HC1-4). E. Western blot of FACS-sorted CD3+ CD8+ T cells from A1, heterozygote parent (A0), and three healthy controls (HC1-3). F. Western blot of FACS-sorted CD3+ CD4+ T cells from A1, heterozygote parent (A0), and three healthy controls (HC1-3). G. Western blot of fetal thymuses from Kindred D (D1-D3) and five fetal thymic controls from 25 weeks old (FT3-FT4) and 31 weeks old (FT1, FT2, FT5). A-G, loading control: Actin.

### The L77P IL-2Rβ missense mutation causes loss of surface expression and function in T cells

At baseline, IL-2Rβ is normally highly expressed on the surface of unstimulated control CD3-CD56+ NKs; however, patients with the L77P mutation have markedly decreased surface expression of IL-2Rβ on NKs, CD4 T cells, and CD8 T cells as assessed by flow cytometry (Figure 2C and data not shown). A healthy heterozygous parent showed intermediate surface expression of IL-2Rβ (Figure 2C). Despite diminished cell surface IL-2Rβ expression, immunoblotting of cytosolic lysates of patient NKs, CD4 T, and CD8 T cells revealed strikingly more IL-2Rβ than healthy controls (Figure 2B-D). This implied that the mutant L77P IL-2Rβ protein was sequestered intracellularly due to misfolding and an inability to properly traffic to the cell surface for subsequent turnover. In keeping with this hypothesis, the faster migration of L77P IL-2Rβ protein relative to WT IL-2Rβ is likely due to incomplete glycosylation branching modifications that are added post-translationally outside the endoplasmic reticulum (ER) (Figure 2D). In addition, the affected neonate (D1) and fetuses (D2, D3) from Kindred D with the more severe p.Q96* stop gain mutation, resulting in a significant truncation, had no IL-2Rβ protein accumulation (Figure 2E).

### Reduced signaling by mutant IL-2Rβ proteins encoded by patient alleles

We reconstituted the IL-2R complex in HEK293T cells via transfection of expression plasmids encoding IL-2Rβ, and IL-2Rγ, JAK-3, and STAT5; this system can transduce a signal from IL-2 to intracellular mediators such as the STAT signaling proteins (John et al. 1999; Majri et al. 2017). We used this system to compare the protein-coding sequences of the wild-type or L77P mutant IL-2Rβ and GFP separated by a P2A sequence under the control of a tetracycline (Tet)-inducible promoter (pHTC). As expected, cells transfected with the wild-type plasmid showed increasing IL-2Rβ surface expression with increasing GFP expression after Tet induction (Figure 3A). However, cells transfected with the pHTC-mutIL2RB plasmid showed no change in IL-2Rβ surface expression with increasing GFP expression, except for very high levels of GFP expression (Figure 3A). Given similar levels of expression of the BFP control for tetracycline-inducible system and GFP expression, wild-type IL-2Rβ is expressed in much greater abundance on the cell surface than the L77P mutant (Figure 3B). As observed in the L77P mutant patient lymphocytes, there is an increase in total cytoplasmic IL-2Rβ protein, despite decreased surface expression, in cells transfected with the mutant (Figure 3C). Confocal imaging of the live HEK293T cells transfected with KDEL-BFP and wtIL-2RB-GFP or mutIL-2RB-GFP indicates that mutIL-2RB-GFP co-localizes with KDEL-BFP and indicating that it is being sequestered in the ER (Figure 3D), as we hypothesized from the patient data. Together these experiments demonstrate that even when the L77P IL-2Rβ is reconstituted in an exogenous HEK293T cell line, the allele encodes a mutant protein that is sequestered in the ER and fails to reach the cell surface, thus recapitulating the patients’ cellular phenotype.

**Figure 3.**
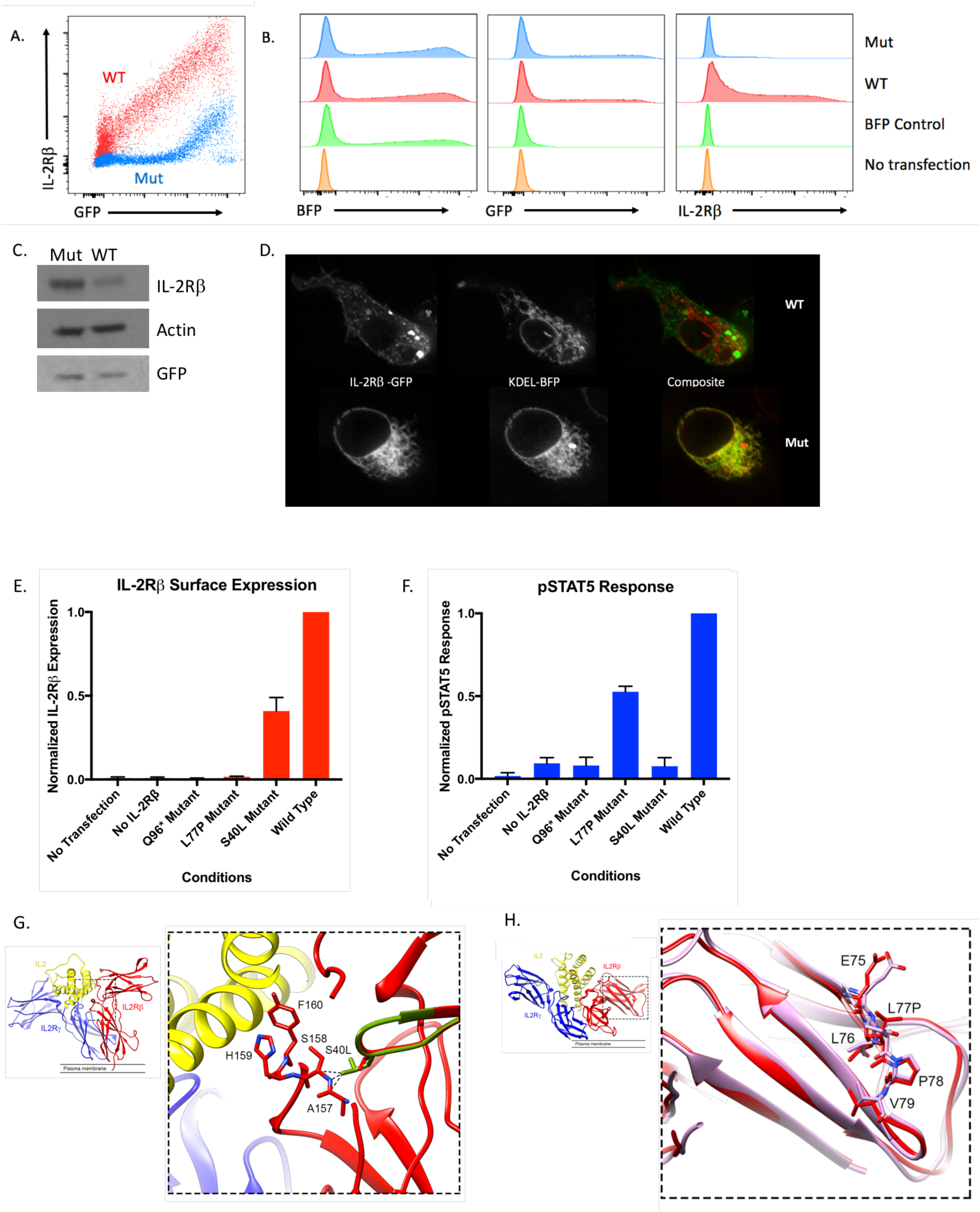
Investigation of IL-2Rβ deficiency mechanisms in a HEK293T transfection model. A. FACS plot of GFP and IL-2Rβ expression by HEK293T cells transfected with pHTC-wtIL2RB (red) and pHR-TetON-BFPor transfected with pHTC-mutIL2RB (blue) and pHR-TetON-BFP. B. Histograms of BFP, GFP, or IL-2RB expression given the listed four transfection conditions: wild-type, mutant, TetON only, and no transfection. C. Western blot of HEK293T cells transfected with pHTC-wtIL2RB-GFP or pHTC-mutIL2RB-GFP. Loading controls: actin and GFP. D. Confocal images of live HEK293T cells co-transfected with KDEL-BFP (ER localization marker) and WT- IL2RB-GFP or Mut-IL2RB-GFP. E. Graph of normalized surface IL-2RB expression in HEK293T cells with exogenous IL-2 receptor system for the three disease-causing IL-2RB mutations. F. Graph of pSTAT5 response to high dose IL-2 in HEK293T cells with exogenous IL-2 receptor system.G. Molecular modeling of the receptor cytokine binding interface. The IL-2Rγ subunit is coloured in blue, IL-2Rβ in red and IL-2 in yellow (PDB: 5M5E). The WT protein (red) is shown with the modelled structure of the S40L variant (green) shown superimposed. The leucine side chain clashes with main chain atoms in the BC2 loop (residues 157-165) of the D2 domain, which contributes directly to IL-2 binding. H. MD simulation of WT IL-2Rβ and the L77P variant, coloured as in panel G. The structure of WT IL-2Rβ after 100 ns of molecular dynamics (MD) simulation (red) is shown superimposed on the structure of the L77P variant after 100 ns MD simulation (pink). The WT protein has β-strand secondary structure at the site of the mutation; the β-stand cannot form with a proline at position 77. The backbone trajectories are shown in semi-transparent color.

Using our reconstituted receptor system, we also compared the Q96* and S40L alleles to the L77P allele for IL-2Rβ surface expression and phosphorylation of STAT5 (pSTAT5) after IL-2 stimulation (Figure 3E-F). As expected, the Q96* allele, which encodes an early stop codon and truncation of IL-2Rβ prior to the transmembrane domain, generates no IL-2Rβ surface expression and shows no pSTAT5 response to IL-2 stimulation (Figure 3E-F). By contrast, the S40L IL2RB allele promoted IL-2Rβ surface expression but had no response to IL-2 stimulation (Figure 3E-F). Molecular modelling of the S40L mutant showed that the substitution introduces steric clashes with main chain atoms in the BC2 loop (residues 157-165) in the D2 domain, which we predict would disrupt the IL-2 binding interface of IL-2Rβ (Figure 3G), consistent with this variant’s lack of responsiveness to IL-2 (Figure 3F). In addition, we performed molecular dynamics (MD) simulations on WT IL-2Rβ and the L77P mutant. After 100 ns of simulation, Pro77 and its two flanking residues adopted a slightly different backbone conformation due to the proline being incompatible with the β-strand, secondary structure adopted by these residues in the WT protein (Figure 3H and S2). Consequently, residues 76-78 in the L77P mutant do not contribute a β-strand to one of the β-sheets in the D1 domain as in WT (Figure 3H). This suggests that the fold of IL-2Rβ D1 is destabilized by the L77P mutation, which is consistent with our functional evidence that the L77P mutant is misfolded and sequestered in the ER. Thus, by using this reconstituted system, we define three distinct mechanisms in humans for IL-2Rβ deficiency by showing that it can occur due to an absence of IL-2Rβ (Q96*), impaired surface expression (L77P), and decreased binding of IL-2 (S40L).

### Patient cells with different mutant alleles have selective signaling defects

We next defined the IL-2 signaling defects in the patient cells by measuring downstream STAT3 and STAT5 phosphorylation in response to high dose IL-2 stimulation (Figure 4). T cells stimulated with high dose IL-2 trigger tyrosine phosphorylation of the cytoplasmic tails of the

**Figure 4.**
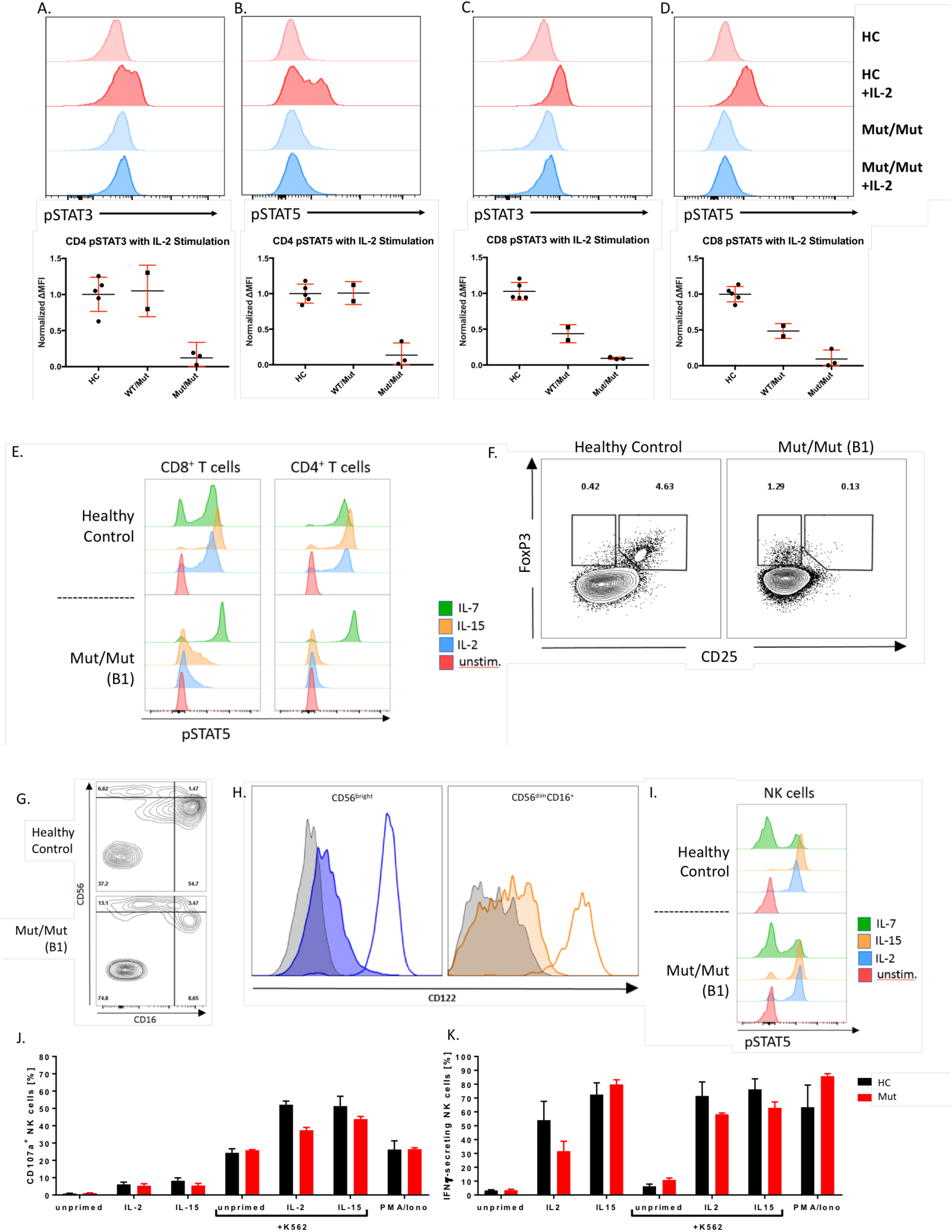
IL-2Rβ deficiency abrogates IL-2 induced STAT3 and STAT5 phosphorylation in peripheral T cells while NK cells retain IL-2/IL-15 responsiveness and effector function. A. Flow cytometry-based measure of STAT3 phosphorylation in CD3^+^ CD4^+^ T cells from healthy controls (HC), heterozygote parent (WT/Mut), and homozygous affected(Mut/Mut). B. STAT5 phosphorylation in CD3^+^ CD4^+^ T cells. C. STAT3 phosphorylation in CD8 T cells. D. STAT5 phosphorylation in CD8^+^ T cells. (red=representative healthy control, blue=representative affected, lighter shade=unstimulated, darker shade=stimulated with 1000U IL-2). E. STAT5 phosphorylation in CD4^+^ and CD8^+^ T cells in response to IL-2, IL-7, and IL-15 stimulation. F. Flow cytometry plot of CD25 and FoxP3 expression in healthy control and homozygous affected. G. Flow cytometry plot of CD16 and CD56 expression in CD3^−^ CD19^−^ lymphocytes in Kindred B. H. FACS plot of IL-2RB expression CD56^bright^ and CD56^dim^ CD16^+^ cell subsets. Gray = isotype control, line only = healthy control, shaded color = Mut/Mut (B1). I. STAT5 phosphorylation in NK cells in response to IL-2, IL-7, and IL-15 stimulation. J. Graph of control and patient NK degranulation when co-cultured with K562 cells and in response to IL-2 and IL-15 stimuli. K. Graph of control and patient NK interferon-gamma (IFN-γ) release when co-cultured with K562 cells and in response to IL-2 and IL-15 stimuli.

IL-2Rβ and IL-2Rγand downstream STAT1, STAT3, and STAT5 phosphorylation via JAK1 and JAK3. Consistent with a loss of function phenotype, we found that CD4+ and CD8+ T cells from patients A1 and B1 failed to phosphorylate STAT5 in response to IL-2 or IL-15 stimulation whereas robust phosphorylation was observed in cells from healthy controls (Figure 4A-E). Control experiments with IL-7 stimulation, which does not signal through IL-2Rβ, showed normal responses indicating that patient T cells were alive and capable of signaling (Figure 4E). Interestingly, CD4+ T cells from A0, the father with a heterozygous L77P genotype, have enough surface expression of IL2-Rβ to phosphorylate STAT3 and STAT5 at a comparable level to healthy controls (Figures 4A and B). By contrast, heterozygous CD8+ T cells cannot offset the decreased IL2-Rβ surface expression leading to a proportional decrease of STAT3 and STAT5 phosphorylation (Figures 4C and D). Thus, surface IL-2Rβ deficiency abrogates downstream STAT phosphorylation in response to IL-2 stimulation in a cell-type- and receptor expression-dependent manner.

In keeping with current understanding of the critical role of IL-2 signaling in the maintenance of regulatory T (Treg) cells in the periphery, the CD25^hi^FoxP3+ CD4+ T cell compartment was almost empty (Figure 4F). Taken together, the profound reduction of STAT5 signalling within the CD4+ T cell compartment and the absence of CD25^hi^FOXP3+ T_reg_s closely mirrors the situation in IL2RB-knock out mice and other known human Treg disorders such as deficiency states of FOXP3 and CD25. Therefore, this could be sufficient to explain the various autoimmune manifestations we observed early in life.

### Hypomorphic nature of L77P IL2RB mutation in NK cells

The NK compartment of IL2RB-knockout mice is almost completely depleted, but our patients bearing hypomorphic mutations instead showed an expansion of NK cells (Supplementary Table 2) and an increase in CD56^bright^ relative to CD56^dim^ NKs (Figure 4G) (Suzuki et al. 1995). Indeed, residual expression of IL-2Rβ^L77P^ was clearly detectable in both NK subsets (Figure 4H), just as it had been on the surface of transfected 293T cells. Moreover, this residual IL-2Rβ expression could sustain IL-2 and IL-15 signal transduction and downstream STAT5 phosphorylation (Figure 4I). The patient’s NK cells also showed comparable effector functions, in terms of interferon gamma (IFN-γ) release and degranulation, relative to healthy controls (Figure 4J and 4K). These data support the conclusion that L77P is a hypomorphic mutation of IL-2Rβ that all but abolishes IL-2 signaling in T cells but still transduces residual signaling in high IL-2Rβ expressing cell subsets like NK cells. As a result, NK cells persist and can respond to the IL-2 and IL-15 that we speculate are normally produced but not consumed by IL-2Rβ-deficient T cells. Moreover, we have preliminary evidence to suggest that certain NK, CD8, and CD4 T cell subsets are more perturbed than others due to variable levels of IL-2Rβ expression (Figure S2). Importantly, we observed an absence of terminally differentiated populations of NKs and CD8 T cells (Figure S2), which may contribute to CMV persistence and autoimmunity. This demonstrates in human cells that mutant IL2RB alleles may confer different levels of impairment in different immune cell types.

## Discussion

Here we describe the first report of autosomal recessive IL-2Rβ deficiency in four pedigrees harboring five affected liveborn children with immunodeficiency and autoimmunity and three perinatally affected fatalities. Our identification of human IL-2Rβ deficiency as a monogenic cause of immunodeficiency and autoimmunity provides insight into one of the principal signaling pathways of the immune response and should prompt prenatal screening of IL-2RB mutations and genetic counseling in at risk families. Clinical hallmarks of the disease include prominent immune dysregulatory phenomena such as enteropathy, skin abnormalities, autoimmune hemolytic anemia, and hypergammaglobulinemia, together with susceptibility to respiratory and herpesvirus infections. We demonstrate that the three mutant alleles cause IL-2Rβ deficiency by different biochemical mechanisms. Kindreds A and B have the hypomorphic L77P IL-2Rβ mutation which interferes with egress from the endoplasmic reticulum. We discovered that this abrogates surface expression and IL-2 signaling in T cells, but that NKs not only retain modest surface expression and responsiveness to IL-2 but quite potent cytolytic activity. Kindred C possesses the S40L IL-2Rβ mutant, which has decreased responsiveness to IL-2 despite being expressed on the cell surface. Our analysis shows that this is due to an amino acid side group clash in the receptor: ligand interaction site. Kindred D has the most severe Q96* IL-2Rβ stop gain mutation. The severity of the mutation is reflected not only the clinical phenotype in the neonate and fetuses but also by the complete absence of IL-2Rβ expression and IL-2 signaling. Despite the differences in mechanism, all the mutations cause IL-2Rβ dysfunction in some manner and lead to a similar constellation of clinical features.

Specifically, the hypomorphic L77P IL-2Rβ mutation highlights the significance of variable IL-2Rβ expression in different lymphocyte subsets as a means of modulating immune function. The L77P mutation causes ER sequestration and thus minimal IL-2Rβ surface expression in patient lymphocytes despite increased total IL-2Rβ protein. This decreased IL-2Rβ surface expression prevents downstream STAT3 and STAT5 phosphorylation following IL-2 stimulation in T cells. However, NK cells are still capable of responding to IL-2 and maintain normal effector function, likely due to the cell’s intrinsic high expression of IL-2RB.

While the human IL-2RB deficiency shares many similarities with the IL2RB knockout (KO) mouse and FoxP3 deficient IPEX patients, there are interesting key differences. Like the knockout mouse (Suzuki et al. 1995), the IL-2RB deficient patients have autoimmune hemolytic anemia, elevated autoantibodies, and hypergammaglobulinemia (IgG and IgE), lymphadenopathy, and splenomegaly. In vitro both the IL-2RB KO mouse and IL-2RB deficient human T cells do not proliferate in response to IL-2 and TCR stimulation. Human IL-2RB disease reveals that deficient IL-2RB also leads to skin abnormalities and enteropathy, which is not seen in the KO mouse. In addition, in the human patients, we observed an expansion of NKs, while a reduction of NKs was recorded in the KO mouse (Suzuki et al. 1997). These observations suggest differences in the regulation and role of IL-2/15Rβ in human and mouse NK maturation (Renoux et al. 2015) and tissue resident memory T cells. In parallel to the KO mouse, the IL-2Rβ deficient patients also lack CD25+ FoxP3+ regulatory T cells and, thus, share many clinical features with IPEX syndrome. As expected, there is significant overlap in the immune-mediated symptoms (enteropathy, dermatitis, and hemolytic anemia) of both IL-2Rβ and FoxP3 deficiency; however the distinguishing component of IL-2RB deficiency is the presence of recurrent infections in addition to severe autoimmune/inflammatory disease. Moreover, only one IL-2RB patient (A1) presented with any endocrinopathy – a hallmark of IPEX. The presence of both immunodeficiency and autoimmune/inflammatory disease as defining features of IL-2RB disease is consistent with the multi-faceted role of IL-2 signaling biology.

The current definitive treatment for IL-2Rβ deficiency is hematopoietic stem cell transplant. Patient A2 received an allogeneic HSCT and has had complete resolution of her symptoms. However, there are high risks associated with HSCT, as exemplified by patient B1, and the hope is that understanding the pathophysiological mechanism of IL-2Rβ deficiency can guide the development of novel therapeutics. For example, if the clinical phenotype is due to hypomorphic IL-2Rβ deficiency, there may be alternative corrective rescue methods or potential treatment strategies. It is feasible to boost IL-2 IL-2R interaction by IL-2 anti-IL-2 antibody complexes (Boyman et al. 2006), IL-2 superkine (Levin et al. 2012), ortho-IL2 analogs (Sockolosky et al. 2018), and IL-2 Fc fusion proteins (Vazquez-Lombardi et al. 2017) as a potential means of hyper-stimulating residual surface IL-2Rβ. Monoclonal anti-human IL-2 antibody MAB602 (mouse S4B6) in complex with IL-2 was found to selectively promote proliferation of effector T cells, while the antibody clone 5344 (mouse JES61) induced proliferation of T_regs_ (Boyman et al. 2006). Similarly, the H9 IL-2 superkine was engineered to have enhanced binding to IL-2Rβ independent of CD25 (Levin et al. 2012). Another approach to hyper-stimulating the IL-2Rβ mutant would be to develop an orthoIL-2 with specific binding to the mutant (Sockolosky et al. 2018). IL-2-Fc fusion proteins could also potentially rescue the IL-2Rβ mutant by inducing proliferation of CD25 deficient T cells without affecting T_regs_ (Vazquez-Lombardi et al. 2017). While these experimental therapies are still a long way from being used at the bedside, low-dose IL-2 therapy, which is currently in Phase II clinical trials for lupus and approved for higher dose use for cancer treatment, may benefit those with hypomorphic IL-2Rβ deficiency by priming their immune systems.

## Acknowledgements

This work was supported by the Wellcome Trust (Investigator Award 083650/Z/07/Z to KGCS, 207556/Z/17/Z to SH, 101908/Z/13/Z to YM), the Division of Intramural Research, National Institute of Allergy and Infectious Diseases, NIH, Merck, Inc, and the UK National Institute of Health Research Cambridge Biomedical Research Centre and the Sir Jules Thorn Charitable Trust (12/JTA to SH). Z.Z. was supported by the NIH-Oxford-Cambridge Scholarship in Biomedical Research program and the NIH M.D./Ph.D partnership program with Harvard Medical School. F.G. was supported by the Deutsche Forschungsgemeinschaft (GO2955/1-1).

The authors thank John Sowerby, Lixin Zheng, Morgan Similuk, Warren Leonard, and Helen Su for their advice and insight. We thank Daniil Prigozhin for advice on molecular dynamics simulations and Patricia Fergelot at the Genome Transcriptome Facility of Bordeaux BIOGECO, INRA for their support in whole exome sequencing. Finally, we thank all the patients described in this manuscript and their families for facilitating this work.

## Methods

### Human Subjects

Written informed consent was provided by all human subjects or their legal guardians in accordance with the 1975 Helsinki principles for enrollment in research protocols that were approved by the Institutional Review Board of the National Institute of Allergy and Infectious Diseases, National Institutes of Health and the Newcastle and North Tyneside Research Ethics Committee 1, UK. Patient and healthy control blood was obtained at Starship Children’s Hospital in Auckland, New Zealand, Addenbrooke’s Hospital in Cambridge, United Kingdom, and Great North Children’s Hospital in Newcastle, United Kingdom under approved protocols.

### Genetic Analysis

DNA was obtained from probands and family members by isolation and purification from peripheral blood mononuclear cells (PBMCs) using Qiagen’s DNeasy Blood and Tissue Kit. The DNA was then submitted for whole exome sequencing (WES) by Illumina sequencers in the United States, United Kingdom, France, and Saudi Arabia. The reads were filtered for sequence quality and then mapped on to the h19 human genome reference by Burrows-Wheeler Aligner with default parameters. Alignment, variant calling, and annotation were performed by the in-house bioinformatics core using the Genome Analysis Toolkit version 3.4 (Broad Institute) and GEMINI (GEnome MINIng). The IL2RB variant was confirmed by Sanger sequencing of PCR amplification products of cDNA, generated by reverse transcription of RNA using SuperScript IV VILO kit (Thermo) and the following PCR primers: F-CCTGTGTCTGGAGCCAAGAT and R-GGGTGACGATGTCAACTGTG (Sigma Aldrich) or F-CCTCACAGTGGTTGGCACA and R-GCACTCTCTCCCTGGGTG (Sigma Aldrich).

### Cells and Media

Primary patient or control PBMCs were obtained from whole blood subjected to Histopaque/Ficoll density gradient separation. The PBMCs were then washed with PBS and frozen in complete RPMI with 10% DMSO in liquid nitrogen for later use or −80°C for transport. HEK293T and K562 cells were obtained from the European Collection of Authenticated Cell Cultures and tested mycoplasma-free (ECACC). Human cells were cultured in RPMI (Sigma Aldrich) or DMEM (Sigmal Aldrich) supplemented with 10% heat-inactivated fetal bovine serum (Sigma Aldrich), 1% penicillin/streptomycin (Gibco), and 1% Glutamax (Gibco). Recombinant human IL-2, IL-7 and IL-15 (Peprotech) was used for stimulation. XVIVO 15 media (Lonza) supplemented with 1-10% human AB serum (Sigma Aldrich) was used for STAT phosphorylation assays.

### Antibodies

The following monoclonal primary rabbit anti-human antibodies from Cell Signaling Technologies (CST) were used for Western blot analysis: anti-IL2RB, anti-GFP, anti-vinculin, and anti-IL2RA. Rabbit anti-beta actin (Abcam) and goat anti-IL2RG (Thermo Fisher Scientific) were also used. Secondary HRP-linked anti-rabbit IgG and anti-goat IgG antibodies (CST) were used to conjugate to the respective primary antibodies. The following flow cytometry antibodies are from Biolegend: CD3-AF700, CD3-PerCp-Cy5.5, CD3-BV705, CD4-Pacific Blue, CD56-PE-Cy7, CD122-PE-Dazzle, CD132-APC, CD25-APC-Cy7, CXCR5-FITC, CD45RA-PerCp-Cy5.5, CD127-APC, HLA-DR-Pacific Blue, and Live/Dead-Zombie Aqua, pSTAT3-AF647, CD20 (2H7), PD-1 (EH12.2H7), CD127 (A019D5), TNFα (MAB11), CD56 (5.1H11), CD56 (HCD56), CD16 (3G8), CD19 (HIB19), CD122 (TU27), CD57 (QA17A04), and Perforin (dG9); Thermo Fisher Scientific: CD4-APC-eF780, CD56-APC-eF780, TCRγδ (B1.1), and TCRVα24Jα18 (6B11); BD Bioscience: CD25-PE, pSTAT5-AF488, CD4 (SK3), CD3 (UCHT1), CD8 (RPA-T8), CD25 (2A3), CCR7 (3D12), CD45RO (UCHL1), Granzyme B (GB11), IFN-γ (B27), STAT5 (47/Stat5), S6 (N7-548), FoxP3 (259D/C7), CD127 (HIL-7R-M21), CD56 (NCAM16.2), CD28 (CD28.2), CD95 (DX2), CD16 (3G8), CD107a (H4A3), CD69 (FN50), IL-2 (5344.111); Miltenyi: CD132 (REA313); and R&D Systems: NKG2C (134591). Cell trace violet (Thermo Fisher Scientific, MA, USA) was used to label K562 cells. Cell viability was assessed using Zombie NIR, 7-AAD (both from Biolegend, CA, USA) or LIVE/DEAD Fixable Green (Invitrogen, Thermo Fisher Scientific, MA, USA). IFN-γ secretion was detected using the IFN-γ secretion assay-detection kit (Miltenyi Biotec, Bergisch Gladbach, Germany).

### Flow Cytometry

Cells were pelleted by centrifugation and stained with antibodies 1:100 dilution in FACS Buffer (1-2% FBS, 0.05% sodium azide, and 2-5 mM EDTA in PBS) at 4°C for 30-60 minutes. The stained cells were then washed with PBS or FACS buffer, pelleted, and resuspended at ~1×10^6^ cells/ml in FACS Fix Buffer (FACS Buffer with 1% PFA) for flow cytometry analysis (Fortessa, Symphony A5, or FACS Aria Fusion systems). The flow data was analyzed using FlowJo or Treestar.

### Western Blot

Cells were lysed with NuPage LDS sample buffer (Thermo Fisher Scientific) at the concentration of 10^5^ cells per 15uL LDS supplemented with 10% BME and Benzonase Nuclease (Sigma Aldrich). The samples were then denatured at 70°C. Protein lysates were separated by SDS-PAGE on 4-12% Bis-Tris precase gels (Invitrogen) and transferred to a PVDF membrane (Invitrogen) by iBlot (Thermo Fisher Scientific) or wet transfer. Membranes were then blocked in milk with 5% Tris-buffered saline with 0.01% Tween-20) TBST for an hour at room temperature and then incubated with primary antibody in milk or 5% BSA overnight at 4°C. The membrane was washed for 3 x 10 minutes with TBST at room temperature and then stained with HRP-linked secondary antibody in milk for 1 hour at room temperature. After 3 x 10 minute washes with TBST and 1 x 10 minute wash with PBS, the membrane was exposed to enhanced chemiluminescent (ECL) substrates (Thermo Fisher Scientific) and developed by film.

### Flow Cytometry Based STAT Phosphorylation Assay

At the NIH, PBMCs were thawed in XVIVO media (Lonza) with 10% human AB serum (Sigma), pelleted, washed with XVIVO, and resuspended in XVIVO media with 1% human AB serum at the concentration of 10^6^ cells/mL. Then the cells were stimulated with 1000U IL-2 (Peprotech) for 10 minutes at 37°C, fixed with BD Fix/Lyse buffer (BD Bioscience) for 10 minutes at 37°C, and then washed with cold PBS with 0.2% BSA. Next, the fixed cells were permeabilized with −20°C methanol for 20 minutes on ice, washed 5 times with cold PBS with 0.2% BSA, and then stained with surface and intracellular flow cytometry antibodies for 30 minutes at 4°C The fixed, permeabilized, and stained cells were washed with PBS and resuspended in PBS with 0.2% BSA for flow cytometry analysis. In Newcastle, thawed PMBCs were rested for 4 hours in serum-free RPMI-media. After the addition of surface markers and a fixable viability dye, 2x 10^5^ cells were stimulated for 10 minutes at 37°C with 100 ng/mL of either IL-2, IL-7, IL-15 or left unstimulated. The Transcription Factor Phospho Buffer set (BD Biosciences) was used to fix and permeabilize cells according to the manufacturer’s instructions. Cells were stained with the remaining surface as well as intracellular markers for 45 minutes at 4°C before cells were washed in TFP Perm/Wash buffer and finally resuspended in FACS buffer for acquisition.

### Site-Directed Mutagenesis

The wild-type pME18S-IL2RB template plasmid (~5000 bp) was obtained from the NIH. Site-directed mutagenesis of T230C (p.L77P) was performed using the In-Fusion HD Cloning Kit (Takara Clontech) and following PCR primers (Sigma Aldrich): F: AGCTGCCCCCCGTGAGTCAA and R: TCACGGGGGGCAGCTCACAGGTTT. The linearized vector was generated by PCR using the CloneAmp HiFi PCR master mix (Takara ClonTech), plasmid template, and primers with the following thermocycling conditions: 35 cycles of 10 seconds at 98°C, 5 seconds at 55°C, and 25s at 72°C. The PCR products were separated on a 1% agarose gel by gel electrophoresis and the desired mutagenized product band was cut out. The PCR product was purified using the NucleoSpin Gel and PCR Clean Up (Takara) from the InFusion Cloning Kit. The linearized, mutagenized product was ligated using the InFusion Enzyme (Takara) to generate the L77P mutant pME18S-IL2RB plasmid. Stellar cells (Takara) were transformed with the new plasmid by heat shock; the transformed cells were plated on ampicillin plates and incubated overnight at 37°C. Plasmid was extracted from individual colonies using the QIAprep Spin MiniPrep Kit (Qiagen). The mutation was confirmed by Sanger sequencing.

### Cloning

Using wild-type and mutant pME18S-IL2RB plasmids as the template, wtIL2RB and mutIL2RB PCR products with AsiSI and SpeI restriction sites were generated using the following primers: F: tagtaggcgatcgcgccaccATGGCGGCCCCTGCTCTGTC and R: ctactaactagtCACCAAGTGAGTTGGGTCCTGAC. The PCR products were purified by gel electrophoresis. Next the gel purified PCR products and pHTC-P2A plasmid (provided by John James) were digested with AsiSI and SpeI restriction enzymes in CutSmart Buffer (NEB) for 2 hours at 37°C and then purified by gel electrophoresis. IL2RB wt and mutant were ligated into separate pHTC-P2A vectors using T4 DNA ligase (NEB). DH5alpha competent bacteria (NEB) were transformed with pHTC-wtIL2RB and pHTC-mutIL2RB and plated on Amp plates overnight. Individual colonies were Sanger sequenced to confirm successful cloning. pHTC-wtIL2RB, pHTC-mutIL2RB, and pGFP (provided by John James) were digested with mLuI and BamHI in NEB3.1 buffer and then purified by gel electrophoresis. Similar to above, GFP was ligated in to the pHTC vectors to generate pHTC-wtIL2RB-P2A-GFP and pHTC-mutIL2RB-P2A-GFP. The final plasmids were transformed in to DH5alpha bacteria, and individual colonies were Sanger sequenced again.

### HEK293T Transfection and Confocal Imaging

HEK293T cells were cultured in complete DMEM or RPMI at 37°C in T75 flasks. 4×10^5^ cells in 2mL media were seeded into 6 well plates and grown overnight at 37°C At 40-50% confluence, the cells were transfected using 97uL OPTI-MEM (Gibco) and 3uL GeneJuice Transfection Reagent (VWR) per 1ug DNA. Cells were transfected with 1:pHTC-wtIL2RB-P2A-GFP and pHR-TetON-P2A-BFP, 2:pHTC-mutIL2RB-P2A-GFP and pHR-TetON-P2A-BFP, 3:pHTC-wtIL2RB-P2A-GFP, 4:pHTC-mutIL2RB-P2A-GFP, and 5:pHR-TetON-P2A-BFP. Six hours after transfection with pHTC-IL2RB-P2A-GFP and pHR-TetON-P2A-BFP, cells were dosed with doxycycline (1ug/ml). The transfected cells were cultured overnight at 37°C, pelleted, washed with PBS, and stained with CD122-PE-Dazzle antibody for flow cytometry analysis. Similarly HEK293T cells were transfected with pHR-wtIL2RB-GFP or pHR-mutIL2RB-GFP and pBFP-KDEL in the same conditions in fibronectin-coated dishes for confocal imaging. An Andor spinning disc confocal microscope system was used to image the live cells at 37°C. Under the same conditions, HEK293T cells were also transfected with pME-IL2RG, pME-JAK3, pME-STAT5-HA, pBFP, and different IL2RB plasmids to reconstitute the IL-2 receptor. After successful transfection, the cells were stimulated with high dose IL-2 and STAT phosphorylation was measured by flow cytometry as described above.

### NK degranulation and interferon gamma release assays

PBMCs were seeded at 2 x 10^5^ per well in a 96-well plate and primed with either IL-2 or IL-15 (100ng/ml each) for 12 hours or left unprimed. After the priming period, cells were coincubated with K562 target cells (E:T ratio of 20:1) for 3 hours. Alongside with K562 exposure the CD107a-antibody was added to the wells. PHA or PMA/Ionomycin were used as positive controls in some wells. To assess Interferon-γ secretion cells were harvested, washed, resuspended in complete RPMI medium and incubated for 45 minutes in the presence of an IFN-γ catch antibody (Miltenyi Biotec, Bergisch Gladbach, Germany) at 37 °C. Surface staining including an IFN-γ detection antibody was carried out for 60 minutes on ice. Degranulation was measured by means of CD107a surface expression. The addition of IL-2 or IL-15 was found to not affect the viability of K562 cells.

### Molecular Modeling

Starting models were derived from a crystal structure of IL-2RB in complex with IL2-IL-2RB and IL-2 determined at 2.3 Å resolution (PDB: 5M5E) (Klein et al., 2017). For the S40L variant, the Leu40 side chain was modelled with COOT (Emsley and Cowtan, 2004) without molecular dynamics (MD) simulation. For the L77P variant, the Pro77 side chain was placed in the experimental electron density of Leu77 with COOT while minimizing clashes with surrounding atoms to achieve a favourable initial geometry. The GROMACS software package (Abraham et al., 2015) was used to set up and run MD simulations. The AMBER99SB-ILDN force field (Lindorff-Larsen et al., 2010) and TIP3P water model were used and the structures placed in dodecahedral boxes with 10 Å padding and surrounded with solvent including water and 150 mM NaCl. After steepest-gradient energy minimization, a modified Berendsen thermostat (2 groups, time constant 0.1 picoseconds, temperature 310 K) followed by a Berendsen barostat (isotropic, coupling constant 0.5 picoseconds, reference pressure 1 bar) were coupled to the system over 100 picoseconds. One hundred-nanosecond runs of unrestrained MD trajectories were produced. After removal of periodic boundary condition artefacts, MD runs were visualized and analysed in UCSF Chimera (Pettersen et al., 2004) and bulk statistics extracted using GROMACS analysis routines.

---

**Figure S1.**
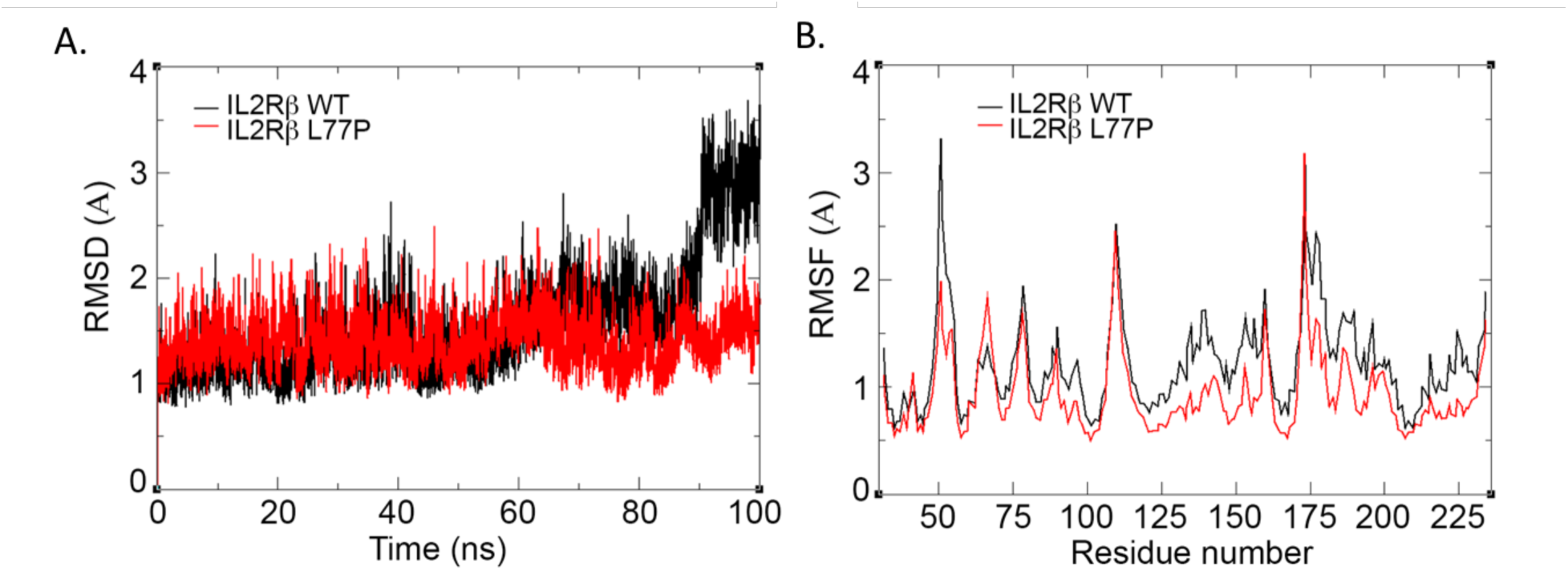
Molecular dynamic simulation of WT and L77P IL-2Rβ structures A. Root mean square deviation (RMSD) between the WT and L77P IL-2Rβ structures over 100 nanoseconds of unrestrained MD simulation with explicit solvent. B. Root mean square fluctuation (RMSF) in WT and L77P IL-2Rβ over the complete trajectory.

**Figure S2.**
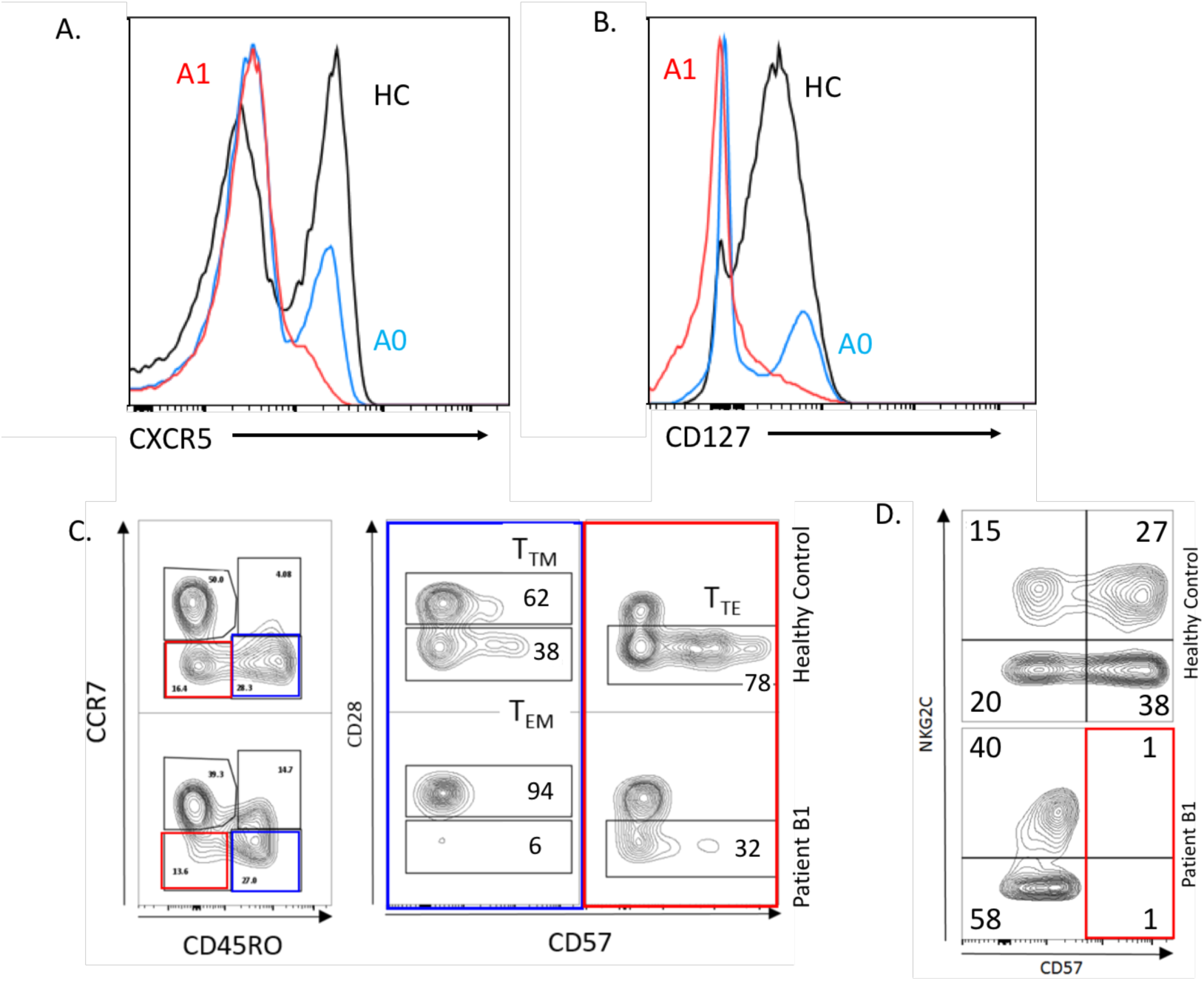
Variable Effect of Hypomorphic L77P IL-2Rβ variant on CD4, CD8, and NK Cell Subsets. Flow cytometry histograms of A. CXCR5 expression in CD25^−^ CD4^+^ T cells in patient A1, heterozygous parent A0, and healthy control (HC) and B. CD127 expression in CD25^−^ CD8^+^ T cells in patient A1, heterozygous parent A0, and healthy control. Flow cytometry contour plots of C. CD28 and CD57 expression of CCR7^−^ CD45RO^+/−^ CD8^+^ T cells in patient B1 and healthy control and D. CD57 and NKG2C expression of CD3-CD56+ NK cells in patient B1 and healthy control.

## References

Abraham, M.J., Murtola, T., Schulz, R., Páll, S., Smith, J.C., Hess, B., and Lindahl, E. GROMACS: High performance molecular simulations through multi-level parallelism from laptops to supercomputers. SoftwareX 2015:1-2, 19–25.

Ahmadzadeh, M. & Rosenberg, S. A. IL-2 administration increases CD4(+)CD25(hi) Foxp3(+) regulatory T cells in cancer patients. Blood 2006; 107: 2409–14.

Boyman, O, Kovar, M, Rubinstein, MP, et al. Selective stimulation of T cell subsets with antibody-cytokine immune complexes. Science 2006; 311: 1924–1927.

Boyman, O, Sprent, J. The role of interleukin-2 during homeostasis and activation of the immune system. Nature Reviews 2012; 12: 180–190.

Busse, D et al. Competing feedback loops shape IL-2 signaling between helper and regulatory T lymphocytes in cellular microenvironments. Proc. Natl Acad. Sci. 2010; 107: 3058–3063.

Caudy, A et al. CD25 deficiency causes an immune dysregulation, polyendocrinopathy, enteropathy, X-linked-like syndrome, and defective IL-10 expression from CD4 lymphocytes. Journal of Allergy and Clinical Immunology. 2007; 119: 482–487.

Emsley, P., and Cowtan, K. Coot: model-building tools for molecular graphics. Acta Crystallogr D Biol Crystallogr 2004: 60, 2126–2132.

Fontenot et al. A function for interleukin 2 in Foxp3-expressing regulatory T cells. Nature Immunol. 2005; 6: 1142–1151.

Hatakeyema, M et al. Interleukin-2 receptor beta chain gene: generation of three receptor forms by cloned human alpha and beta chains cDNA. Science 1989; 1989: 551–556.

Hinks, A et al. Dense genotyping of immune-related disease regions identifies 14 new susceptibility loci for juvenile idiopathic arthritis. Nature Genetics 2013; 45(6): 664–669.

John, S et al. The Significance of Tetramerization in Promoter Recruitment by Stat5. Mol. Cell Biol. 1999; 19(3): 1910–1918.

Klein, C., Waldhauer, I., Nicolini, V.G., Freimoser-Grundschober, A., Nayak, T., Vugts, D.J., Dunn, C., Bolijn, M., Benz, J., Stihle, M., et al. Cergutuzumab amunaleukin (CEA-IL2v), a CEA-targeted IL-2 variant-based immunocytokine for combination cancer immunotherapy: Overcoming limitations of aldesleukin and conventional IL-2-based immunocytokines. Oncoimmunology 2017: 6, e1277306.

Levin, AM et al. Exploiting a natural conformational switch to engineer an interleukin-2 ‘superkine.’ Nature 2012; 24: 352–359.

Liao, W et al. Interleukin-2 at the Crossroads of Effector Responses, Tolerance, and Immunotherapy. Immunity. 2013; 38: 13–25.

Lindorff-Larsen, K., Piana, S., Palmo, K., Maragakis, P., Klepeis, J.L., Dror, R.O., and Shaw, D.E. Improved side-chain torsion potentials for the Amber ff99SB protein force field. Proteins 2010; 78, 1950–1958.

Majri, S et al. STAT5B: A Differential Regulator of the Life and Death of CD4 + Effector Memory T Cells. J Immunol. 2017; 200(1):110–118.

Malek et al. CD4 Regulatory T Cells Prevent Lethal Autoimmunity in IL-2R -Deficient Mice: Implications for the Nonredundant Function of IL-2. Immunity 2002; 17:167–178.

Moffatt, MF et al. A large-scale, consortium-based genomewide association study of asthma. NEJM 2010; 363(13): 1211–1221.

Pettersen, E.F., Goddard, T.D., Huang, C.C., Couch, G.S., Greenblatt, D.M., Meng, E.C., and Ferrin, T.E. UCSF Chimera--a visualization system for exploratory research and analysis. J Comput Chem 2004; 25: 1605–1612.

Renoux, VM et al. Identification of a Human Natural Killer Cell Lineage-Restricted Progenitor in Fetal and Adult Tissues. Immunity 2015; 43: 394–407.

Scharfe, N et al. Human immune disorder arising from mutation of the alpha chain of the interleukin-2 receptor. Proc. Natl Acad. Sci. 1997; 94: 3168–3171.

Sockolosky, JT et al. Selective targeting of engineered T cells using orthogonal IL-2 cytokine receptor complexes. Science 2018; 359:1037–1042.

Suzuki, H et al. Deregulated T cell activation and autoimmunity in mice lacking interleukin-2 receptor beta. Science 1995; 268: 1472–1476.

Suzuki, H et al. Abnormal Development of Intestinal Intraepithelial Lymphocytes and Peripheral Natural Killer Cells in Mice Lacking the IL-2 Receptor Beta Chain. JEM 1997; 185: 499–505.

Takeshita, T et al. Cloning of the gamma chain of the human IL-2 receptor. Science 1992; 257:379–382.

Vazquesz-Lombardi, R et al. Potent antitumor activity ofinterleukin-2-Fc fusion proteins requires Fc-mediated depletion of regulatory T cells. Nat. Commun 2017; 8:15373.

Waldmann, TA. The biology of interleukin-2 and interleukin-15. Nature Rev. Immunol. 2006; 6: 595–601.

Wang, X, Rickert, M, Garcia, KC. Structure of the quaternary complex of interleukin-2 with its a, b, and g receptors. Science 2005; 310:1159–1163.

Willerford, DM et al. Interleukin-2 receptor alpha chain regulates the size and content of the peripheral lymphoid compartment. Immunity 1995; 3:521–530.

Ye, C, Brand, D, Zhong, S. Targeting IL-2: an unexpected effect in treating immunological diseases. Signal Transduction and Targeted Therapy 2018; Online: https://www.nature.com/articles/s41392-017-0002-5#ref-CR77

